# Cleavage Site Specificity for Processing of Farnesylated Prelamin A by the Zinc Metalloprotease ZMPSTE24

**DOI:** 10.1101/2020.08.17.254532

**Authors:** Timothy D. Babatz, Eric D. Spear, Wenxin Xu, Olivia L. Sun, Laiyin Nie, Elisabeth P. Carpenter, Susan Michaelis

## Abstract

The integral membrane zinc metalloprotease ZMPSTE24 is important for human health and longevity. ZMPSTE24 performs a key proteolytic step in maturation of prelamin A, the precursor of the nuclear scaffold protein lamin A. Mutations in the genes encoding either prelamin A or ZMPSTE24 that prevent cleavage cause the premature aging disease Hutchinson Gilford Progeria Syndrome (HGPS) and related progeroid disorders. ZMPSTE24 has a novel structure, with seven transmembrane spans that form a large water-filled membrane chamber whose catalytic site faces the chamber interior. Prelamin A is the only known mammalian substrate for ZMPSTE24, however, the basis of this specificity remains unclear. To define the sequence requirements for ZMPSTE24 cleavage, we mutagenized the eight residues flanking the prelamin A scissile bond (TRSY↓LLGN) to all other 19 amino acids, creating a library of 152 variants. We also replaced these eight residues with sequences derived from putative ZMPSTE24 cleavage sites from amphibian, bird, and fish prelamin A. Cleavage of prelamin A variants was assessed using an *in vivo* yeast assay that provides a sensitive measure of ZMPSTE24 processing efficiency. We found that residues on the C-terminal side of the cleavage site are most sensitive to changes. Consistent with other zinc metalloproteases, including thermolysin, ZMPSTE24 preferred hydrophobic residues at the P1’ position (Leu647), but in addition, showed a similar, albeit muted, pattern at P2’. Our findings begin to define a consensus sequence for ZMPSTE24 that helps to clarify how this physiologically important protease functions and may ultimately lead to identifying additional substrates.

## Introduction

Proteases play key roles in diverse biological processes relevant to human health and disease. Defining their substrate cleavage specificity is essential to fully understand protease physiological function (1,2). ZMPSTE24 is a zinc metalloprotease that is critical for the proteolytic maturation of lamin A, an intermediate filament protein and component of the nuclear lamina (3-9). The lamin A precursor, prelamin A, encoded by the *LMNA* gene, undergoes processing at its C-terminal CAAX motif (C is cysteine, A is usually an aliphatic residue, and X is any amino acid). CAAX processing includes farnesylation of the cysteine residue, endoproteolytic cleavage of the C-terminal tripeptide (-AAX), and carboxyl methylation of the farnesylated cysteine (10-13). Subsequently, ZMPSTE24 proteolytically removes a C-terminal segment of prelamin A, including the farnesylated cysteine, by cleavage between residues Y646 and L647 to produce mature lamin A (3,9,14,15) (Figure 1A).

**Figure 1.**
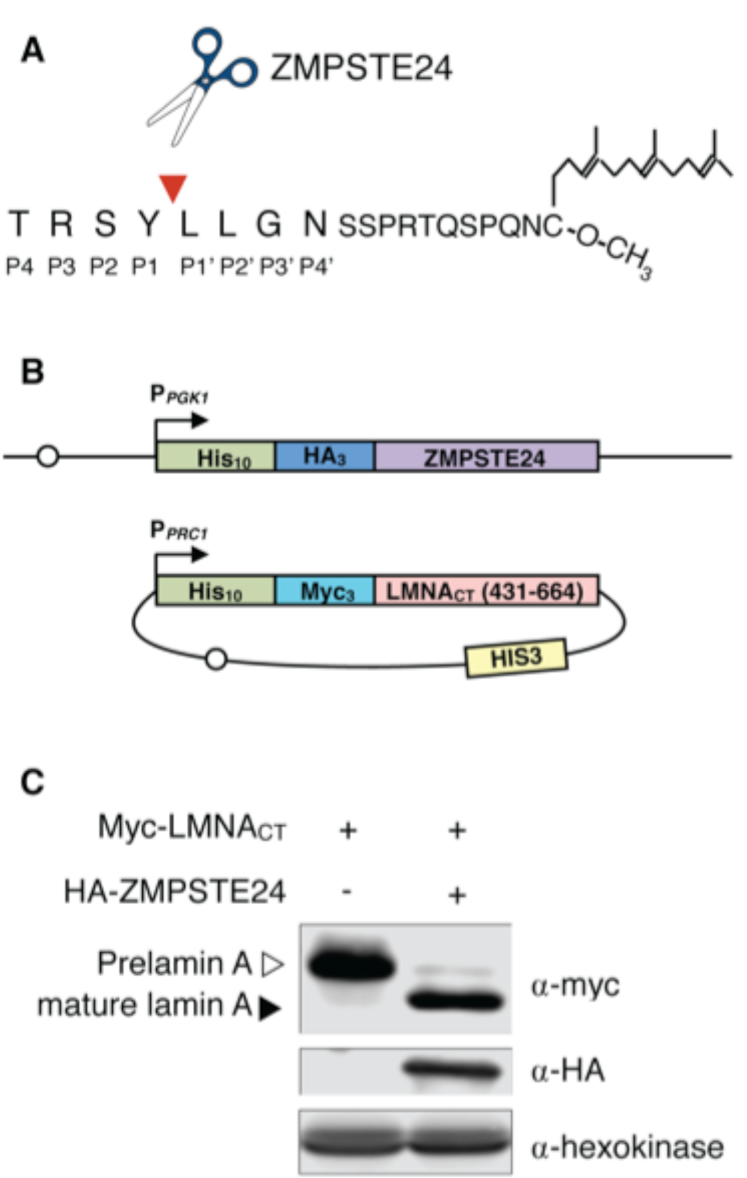
The humanized yeast system (version 2.0) used to assess human ZMPSTE24 cleavage of human prelamin A. (A) Schematic of the C-terminus of prelamin A after farnesylation, CAAX cleavage, and carboxymethylation. The ZMPSTE24 cleavage site between prelamin A residues Y646 and L647 is indicated by a red arrowhead and scissors. Substrate residues mutagenized in this study at positions P4 through P4’ are labeled. (B) Schematic of the humanized yeast system (version 2.0) used to assess processing of mutant forms of prelamin A. Full-length human ZMPSTE24 with an N-terminal 10His-3HA epitope tag is expressed from the PGK1 promoter (P_*PGK1*_) and is chromosomally integrated into a *ste24Δ* strain background SM4826 at the *TRP1* locus, resulting in strain SM6303, which contains 2 tandem copies of the construct shown. The prelamin A substrate on plasmid pSM3393 contains amino acids 431-664 from the C terminus of prelamin A, encoded by human *LMNA* (referred to as LMNA_CT)_, is N-terminally tagged with a 10His-3myc epitope and is expressed from the *PRC1* promoter (P_*PRC1*_) on a centromeric *HIS3* plasmid. This substrate was mutagenized and transformed into strain SM SM6303. Open circles represent centromeric sequences on the chromosome and plasmid. (C) Lysates from yeast strains *ste24Δ* (SM4826; left) and *ste24Δ HA-ZMPSTE24* (SM6303; right) transformed with the prelamin plasmid pSM3393 were analyzed for prelamin A processing by SDS-PAGE and western blotting. Prelamin A and mature lamin A were detected with anti-Myc antibodies; ZMPSTE24 was detected with anti-HA antibodies. Hexokinase serves as a loading control.

Lack of this ZMPSTE24 cleavage event due to mutations in the *LMNA* or *ZMPSTE24* genes results in accumulation of permanently farnesylated and carboxyl methylated prelamin A, which causes progeroid diseases (8,16-18). The premature aging disorder Hutchinson-Gilford Progeria Syndrome (HGPS) is due to a *LMNA* splicing mutation that deletes 50 amino acids from prelamin A, including the ZMPSTE24 processing site (8,19-21). The related progeroid diseases mandibuloacral dysplasia type-B (MAD-B) and restrictive dermopathy (RD) are caused by *ZMPSTE24* mutations (22-26). Disease severity appears to correlate with the extent of ZMPSTE24’s deficiency in cleaving prelamin A (27,28). Some evidence suggests that decreased expression or activity of ZMPSTE24, resulting in the accumulation of prelamin A, may also contribute to normal physiological aging (29).

ZMPSTE24, like its yeast counterpart Ste24, is a multispanning membrane protease with its active site zinc and catalytic residues located inside of a large enclosed water-filled chamber in the membrane created by seven transmembrane spans (30,31). Like other metalloproteases of the gluzincin family, including soluble thermolysin, ZMPSTE24/Ste24 contains an HEXXH motif (31-35). The HEXXH glutamate activates a water molecule for hydrolysis of the scissile bond and the two histidines (along with a glutamate outside of the HEXXH motif) coordinate the zinc atom. ZMPSTE24 is sometimes classified as an intramembrane protease (36), although others argue against this classification since, unlike rhomboids and other intramembrane proteases that cleave substrates within the lipid bilayer, ZMPSTE24/Ste24 substrates are cleaved within the aqueous environment of the membrane-embedded chamber where the catalytic domain resides (34,37). The exact mechanism by which the prelamin A substrate reaches the ZMPSTE24 interior remains unknown; however, the fenestrations observed in the human ZMPSTE24 and yeast Ste24 crystal structures could provide access (30,31). Furthermore, prelamin A is likely too large (∼74 kD) to fit entirely within the catalytic chamber, suggesting that only the farnesylated C-terminal domain of prelamin A must be threaded into the chamber for cleavage.

The substrate specificity and cleavage site determinants for ZMPSTE24/Ste24 have not been well characterized. To date only two specific *in vivo* substrates are known, mammalian prelamin A and yeast **a**-factor, respectively, and both are farnesylated (11,38). Studies *in vivo* and *in vitro* have shown that yeast Ste24 can mediate two cleavage steps for **a**-factor: endoproteolytic removal of the –AAX residues of the CAAX motif CVIA (in which Ste24 plays a redundant role with another protease, Rce1), and a key upstream cleavage between residues T7 and A8 – which Ste24 performs uniquely (39-44). ZMPSTE24 cleaves prelamin A at the upstream site between Y646 and L647, discussed above. While a role for ZMPSTE24 in removal of the –AAX residues of the prelamin A CAAX motif in redundancy with Rce1 has often been suggested, recent evidence suggests that ZMPSTE24 may only poorly cleave the CAAX motif (CSIM) of prelamin A (45). It is notable that both known substrates for ZMPSTE24/Ste24, prelamin A and **a**-factor, are farnesylated. *In vivo*, farnesylation is absolutely required for cleavage of both (14,40,42,46-49). However, *in vitro*, farnesylated prelamin A peptides are cleaved properly, but imperfect cleavage events are also observed for non-farnesylated peptides indicating that farnesyl contributes mainly to fidelity *in vitro* (31,45). Furthermore, with unmodified peptides unrelated to prelamin A, cleavage events can also be detected *in vitro* (50). It is possible that ZMPSTE24/Ste24, which localize to the inner nuclear membrane and endoplasmic reticulum, may cleave substrates in addition to prelamin A and **a**-factor *in vivo*. In both yeast and human cells, clearance of “clogged” ER translocons was shown to require enzymatically active ZMPSTE24/Ste24 suggesting a role in removal of poorly translocating proteins stuck in the Sec61 pore (51,52). In addition, Ste24 was recently implicated in a secretion pathway for proteins that lack a signal sequence (53). ZMPSTE24 also contributes to protecting cells against enveloped viruses, however its role in this process remains enigmatic since catalytic activity appears to be dispensable for this function (54,55).

Apart from the known scissile bond cleaved by ZMPSTE24 in prelamin A (between Y646 and L647) (14,15,56), and by Ste24 in yeast **a**-factor (between T7 and A8) (40,42), there is limited information about the rules governing substrate cleavage. Studies of numerous proteases have employed peptide libraries with variants surrounding the scissile bond, in conjunction with quantitative mass spectrometry to classify the efficiency of various residues for substrate cleavage (57,58). However, the requirement of prelamin A farnesylation for optimal cleavage by ZMPSTE24, along with the difficulty of synthesizing prenylated peptides, and the challenges of purifying and maintaining activity of ZMPSTE24 as a multispanning membrane protein, introduces several technical challenges that make *in vitro* approaches less than ideal. Here, we used a variation of our previously characterized humanized yeast assay (28,59) to define the rules of allowable substitutions at each of the 8 residues flanking the prelamin A cleavage site in order to gain information about how the enzyme works, to assess potential prelamin A disease alleles, as well as to eventually help to identify other putative ZMPSTE24/Ste24 substrates.

## Results

### A humanized yeast system for analysis of ZMPSTE24 cleavage of prelamin A

We previously demonstrated that expression of human ZMPSTE24 together with the C-terminus of human prelamin A in a yeast strain deleted for endogenous *STE24* (*ste24Δ*) provides an ideal model system for studying lamin A maturation *in vivo* (28,59). The initial steps of lamin A biogenesis are mediated by the yeast CAAX processing machinery, resulting in a farnesylated, carboxymethylated prelamin A substrate (Figure 1A). The humanized yeast system used in the present study (called version 2.0) is shown in Fig 1B. It features Myc-tagged prelamin A (residues 431-664) expressed from a plasmid in a yeast *ste24Δ* strain background with chromosomally integrated HA-tagged *ZMPSTE24* (Figure 1B). Western blotting shows that cleavage of the prelamin A substrate is dependent on the presence of ZMPSTE24, and at steady-state, approximately 80-90% of the substrate is in the mature lamin A form (Figure 1C). This version 2.0 humanized yeast system is ideal for maintaining consistent expression of ZMPSTE24, due to its chromosomal integration, while allowing easy introduction by plasmid transduction of a large collection of prelamin A mutants. This contrasts with our previously described system (version 1.0) optimized to study disease alleles of *ZMPSTE24*; in that case *LMNA* was chromosomally integrated and plasmids bore the mutant forms of ZMPSTE24 (28,59).

### Comprehensive Scanning Mutagenesis of the Prelamin A Cleavage Site

The cleavage site specificity for a protease can potentially involve residues both N- and C-terminal to the scissile bond. To methodically survey the importance of residues flanking the prelamin A cleavage site, we mutagenized residues in the P4-P4’ positions shown in Figure 1A to all of the other 19 amino acids. (We use the standard nomenclature of Berger and Schechter (60) in which the unprimed and primed residues refer to the N- and C-terminal sides of the scissile bond respectively, with cleavage occurring between P1 and P1’). This allowed us to assay all 20 possible residues at positions P4-P4’ for cleavage of prelamin A. Plasmids expressing individual prelamin A variants were transformed into the ZMPSTE24-expressing strain SM6303, and the steady-state cleavage efficiency in cells was calculated as a ratio between processed (mature) to total (unprocessed + processed) signal. In all experiments, the wild-type (WT) residue and 19 mutants were assayed in parallel, and cleavage efficiency relative to WT (set to 100%) was determined. Three independently grown replicate cultures were tested for each variant. Results for mutations N-terminal to the cleavage site (residues P1-P4) are shown in Figure 2, and for mutations C-terminal to the cleavage site (residues P1’-P4’) in Figure 3. (See also the heatmap in Figure 4.). This analysis reveals several important insights that are discussed below.

**Figure 2.**
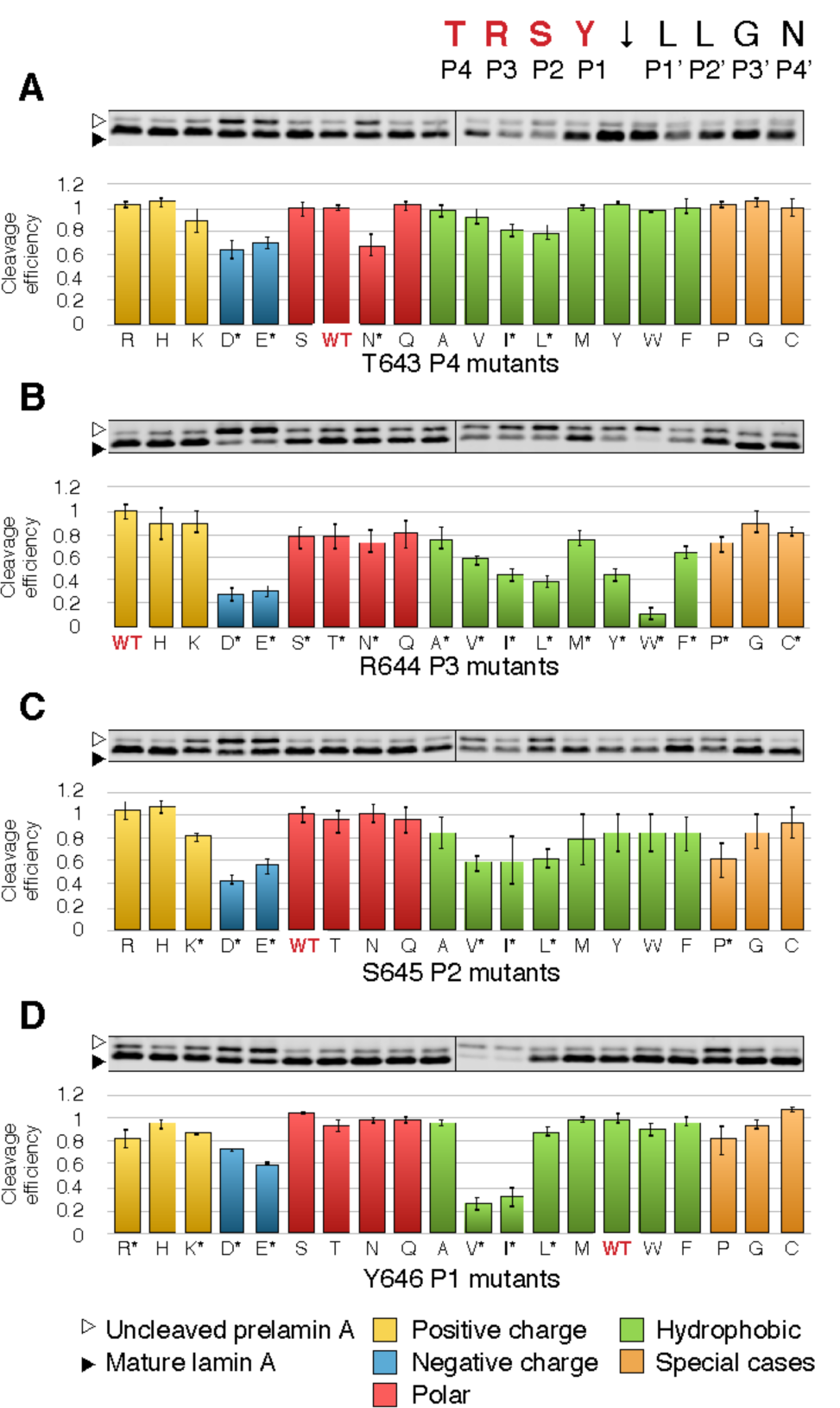
Comprehensive scanning mutagenesis of positions P4—P1 in prelamin A, changing to all other 19 amino acids. Western blots for all possible variants of residues T643 (A), R644 (B), S645 (C), and Y646 (D) are shown. Uncleaved prelamin A (empty arrowhead) and mature lamin A (filled arrowhead) are indicated. For each residue, wild-type (WT) is indicated in red. Residues are color-coded as indicated. Cleavage efficiency is calculated by dividing mature lamin A by total lamin A abundance (cleaved + uncleaved). WT for each residue is normalized to 1 and all mutants are plotted relative to it. Three biological replicates were performed for each mutant. Asterisks by the residue name indicate mutations in which processing is statistically significantly different than WT, Student’s T-test, two-tailed, unpaired, N=3.

**Figure 3:**
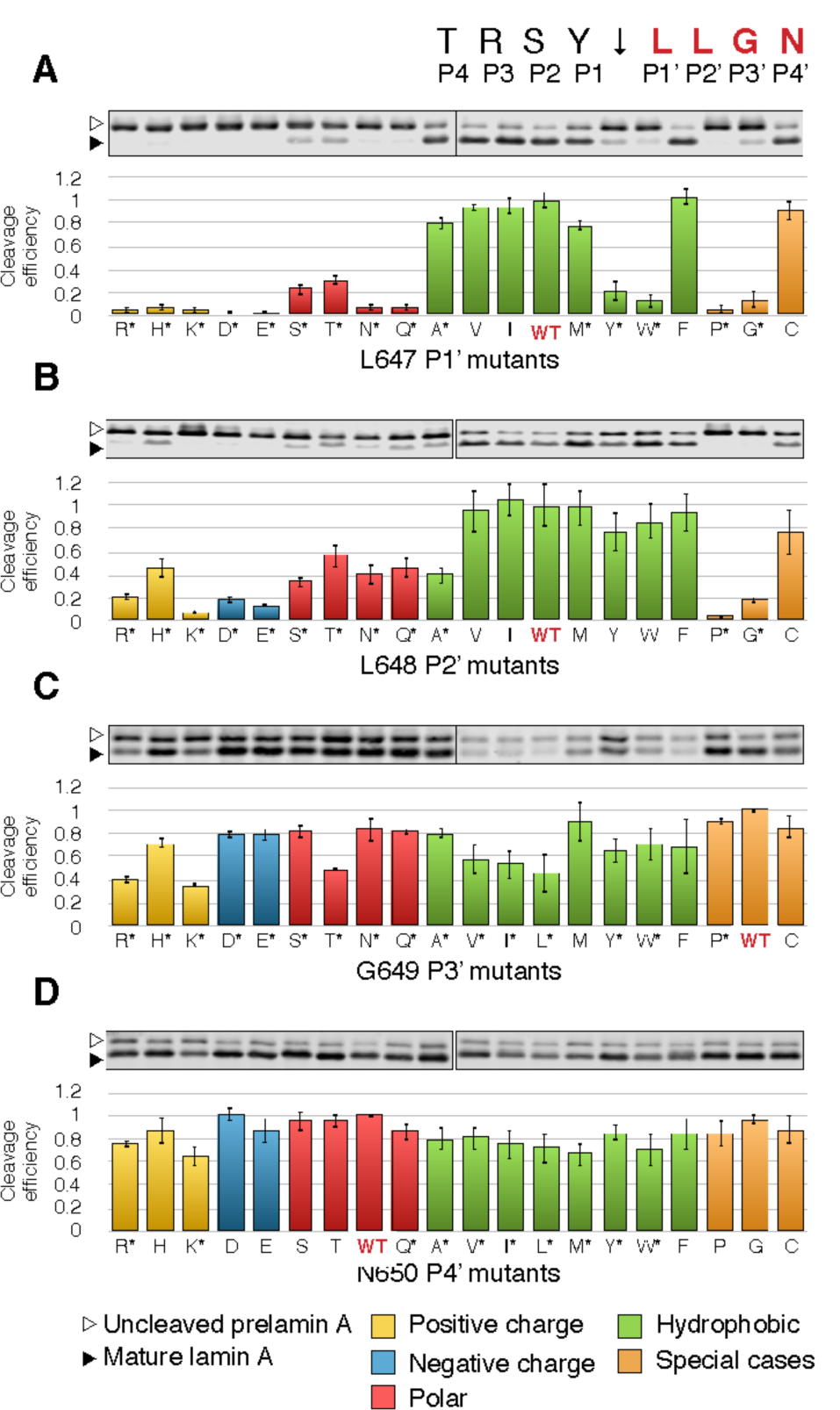
Comprehensive scanning mutagenesis of positions P1’—P4’ in prelamin A to all other 19 amino acids. Western blots for all possible variants of residues L647 (A), L648 (B), G649 (C), and N650 (D), are shown. For each residue, WT is indicated in red. Annotation of uncleaved and mature lamin A, quantification of cleavage, and statistical analysis are performed as explained in the legend for Figure 2.

**Figure 4:**
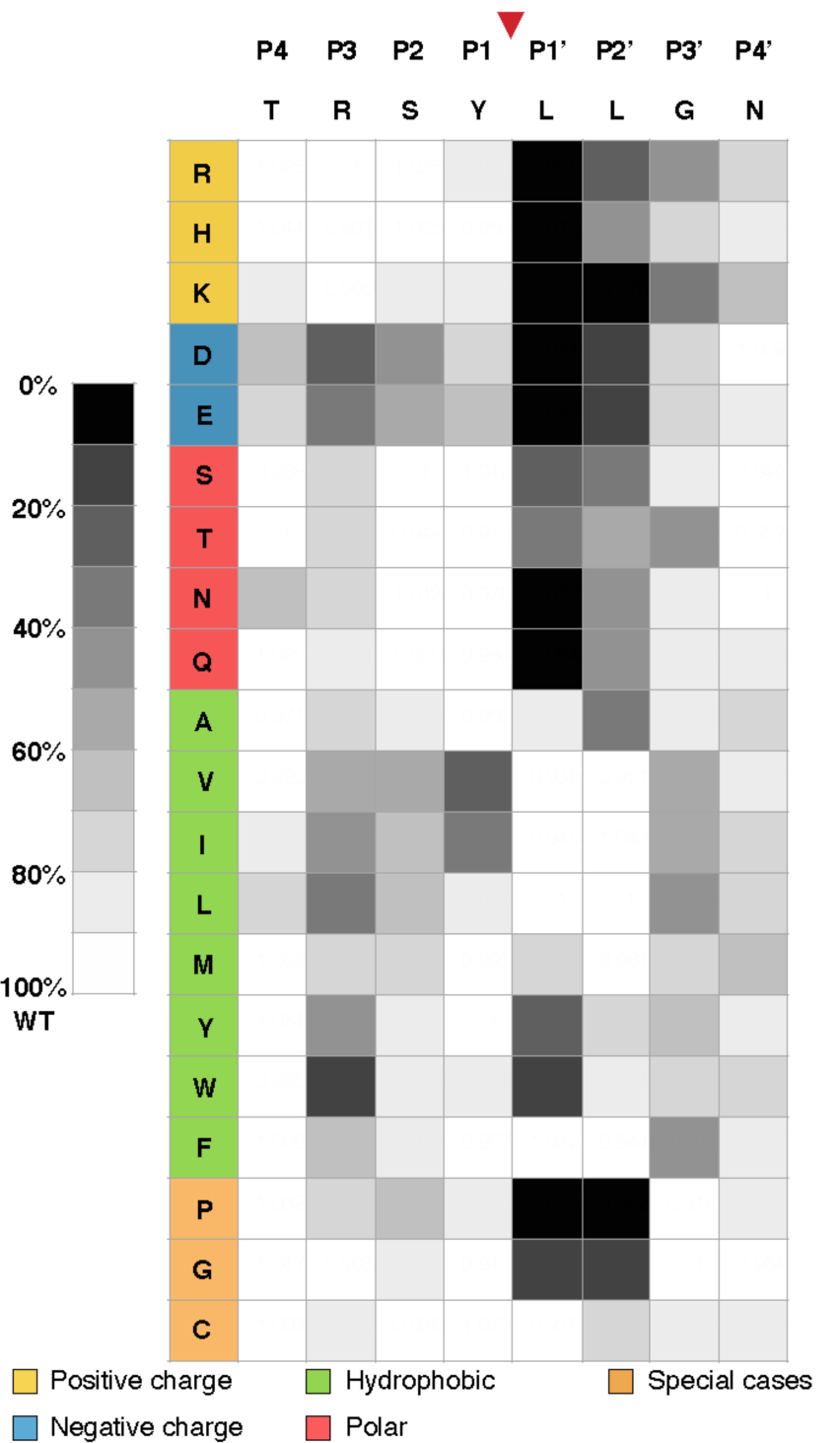
Heatmap showing the ZMPSTE24 cleavage efficiency of prelamin A for every individual amino acid substitution in residues P4-P4’. The P4-P4’ region surrounding the prelamin A cleavage site was comprehensively interrogated by positional scanning mutagenesis, as shown in Figures 2 and 3. The effects of individual amino acid substitutions at each site were scored relative to WT at 100% cleavage efficiency and assigned into ten bins. Residues are shaded in accordance with cleavage efficiency, ranging from WT level of cleavage (white) to poor or no cleavage (black).

For the most distal residues (P4 and P4’), only a few mutants showed effects on cleavage, and none of these were dramatic (Figures 2 and 3). Changes in T643 (the P4 position) to almost any other amino acid had little effect on substrate cleavage (Figure 2A). Even the strongest mutants (T643D, E and N), still retained significant cleavage of at least 60% compared to the WT allele. Similarly, the P4’ position (N650) is also largely insensitive to changes, with no mutants reducing cleavage to less than 60% (Figure 3D). Together, these data suggest the cleavage site recognition motif is generally defined by positions from P3 to P3’.

N-terminal to the scissile bond, at positions P3 to P1, substitutions to aspartate, glutamate, valine, isoleucine, and tryptophan disrupts prelamin A cleavage, showing minor to severe disruptions depending on the residue queried (Figure 2B-D). Just N-terminal to the scissile bond at position P1 (Y646), substitutions that change the aromatic residue tyrosine to the aliphatic residues isoleucine or valine impaired cleavage significantly (>50%), while other substitutions showed mild or no effects. At P2 (S645), only a single substitution to aspartate reduced cleavage to lower than 50%. In contrast, the arginine at position P3 (R644) is the most sensitive to substitution, with the negatively charged residues aspartate and glutamate, the hydrophobic residues isoleucine, leucine, tyrosine, and particularly tryptophan disrupting cleavage significantly (to <30% residual activity; Figure 2B). Many other R644 substitutions also showed effects on cleavage efficiency, albeit modest (<20% reduction of cleavage). This residue is of particular interest because of the association of R644 mutations with diverse laminopathy disease phenotypes (61) and is discussed further below. Taken together only a few scattered substitutions in the P3 to P1 residues N-terminal to the scissile bond of prelamin A showed significant processing defects, but no strong pattern for substrate cleavage requirements in this region was evident.

In contrast to residues N-terminal to the cleavage site, residues C-terminal to the cleavage site at the P1’-P3’ were more sensitive to mutation, especially at the P1’ and P2’ positions, and preference patterns were evident (Figure 3A-C; see also Figure 4). Leucines at positions P1’ and P2’ emerged as critical to ZMPSTE24 processing of prelamin A, with hydrophobic residues strongly preferred at these positions (with the exception of bulky aromatics tyrosine and tryptophan for L647) (Figure 3A and B). Consistent with previous studies (28,48,56), the L647R mutation decreased cleavage efficiency to about 5% relative to WT (Figure 3A). L647R has long been known to disrupt processing and has been reported in a patient with a progeroid disorder (62). Our comprehensive mutagenesis shows that all other charged residues, as well as polar residues, proline, and glycine, nearly abolish processing at L647. L648 shows a very similar pattern, albeit more muted (Figure 3B and Figure 4), with several notable differences. The bulky aromatic hydrophobic residues tyrosine and tryptophan that are disruptive at L647 (P1’) are tolerated at L648 (P2’). Substitution with alanine is tolerated at L647, but disruptive at L648 which had been observed previously in a limited mutagenesis scan in mammalian cells (63). Substitution of proline and glycine at both L647 and L648 largely abolish ZMPSTE24 processing, while these substitutions are well-tolerated at all other positions flanking the cleavage site.

We note that some substitution mutations in both the primed and unprimed positions (*i.e*. Y646V, I and G649V, I, L) showed a low amount of total (unprocessed + processed) signal, possibly due to impaired translatability or RNA stability stemming from poor codon usage or aberrant mRNA structure. However, this did not appear to impact cleavability *per se* and attempts to change several of these substitutions to optimal codons for yeast did not increase the amount or alter extent of cleavage. It is also possible that these versions of prelamin A are degraded; however, we found that proteasome inhibitors did not increase their amounts (data not shown).

Using the cleavage data from Figures 2 and 3, we constructed a heatmap of cleavage efficiency for the prelamin A residues we tested to further help detect patterns (Figure 4). Generally, it is evident from the darker shades in the heatmap that the C-terminal side of the cleavage site (P1’-P3’) is far more sensitive to mutation than the N-terminal side (P3-P1), and that the P4 and P4’ positions show low preference for specific residues. No mutations to the N-terminal side of the cleavage site completely abolish processing, while many C-terminal mutations do. However, the severity of two changes at changes at P1 (Y646V and Y646I) and several at P3 (R644D, E, I, L,Y, W) are notable. As shown for other zinc metalloproteases, a hydrophobic amino acid at the P1’ position is strongly preferred by ZMPSTE24 (with the exception of tyrosine or tryptophan), which fits with the previous suggestion that the proposed S1’ binding pocket in ZMPSTE24 may be a hydrophobic patch (31). In contrast to other metalloproteases, ZMPSTE24 also appears to prefer a hydrophobic residue at P2’ suggesting an extended hydrophobic patch that encompasses both S1’and S2’.

### Multiple Evolutionarily Conserved Substitutions at the Prelamin A Cleavage Site are Tolerated by ZMPSTE24

In the comprehensive mutational analysis above, we could analyze only one amino acid change at a time. Because we were also interested in whether multiple combinatorial amino acid substitutions at the prelamin A cleavage site would be tolerated by ZMPSTE24 we turned to evolutionary information. The *LMNA* gene shows significant conservation throughout vertebrates. We performed a protein sequence alignment for several mammals, birds, fish, and amphibians, which indicated evolutionary conservation of the C-terminus of prelamin A (Figure 5A). The CAAX motif is highly conserved through species, with the farnesylated cysteine invariantly four residues from the C-terminus for all species and a CAAX consensus sequence of C-S-I/V-M. However, there was a marked degree of variation in the residues surrounding the predicted ZMPSTE24 cleavage site, ranging from three to seven amino acid differences from human *LMNA*, depending on the species (Figure 5B, see residues in red). (It should be noted that only human, mouse, and chicken prelamin A have definitively been shown to be processed (3,9,49,56)). Importantly, all species had hydrophobic residues in the critical P1’ and P2’ residues just C-terminal to the cleavage site and generally conform to the rules of the heatmap in Figure 4.

**Figure 5:**
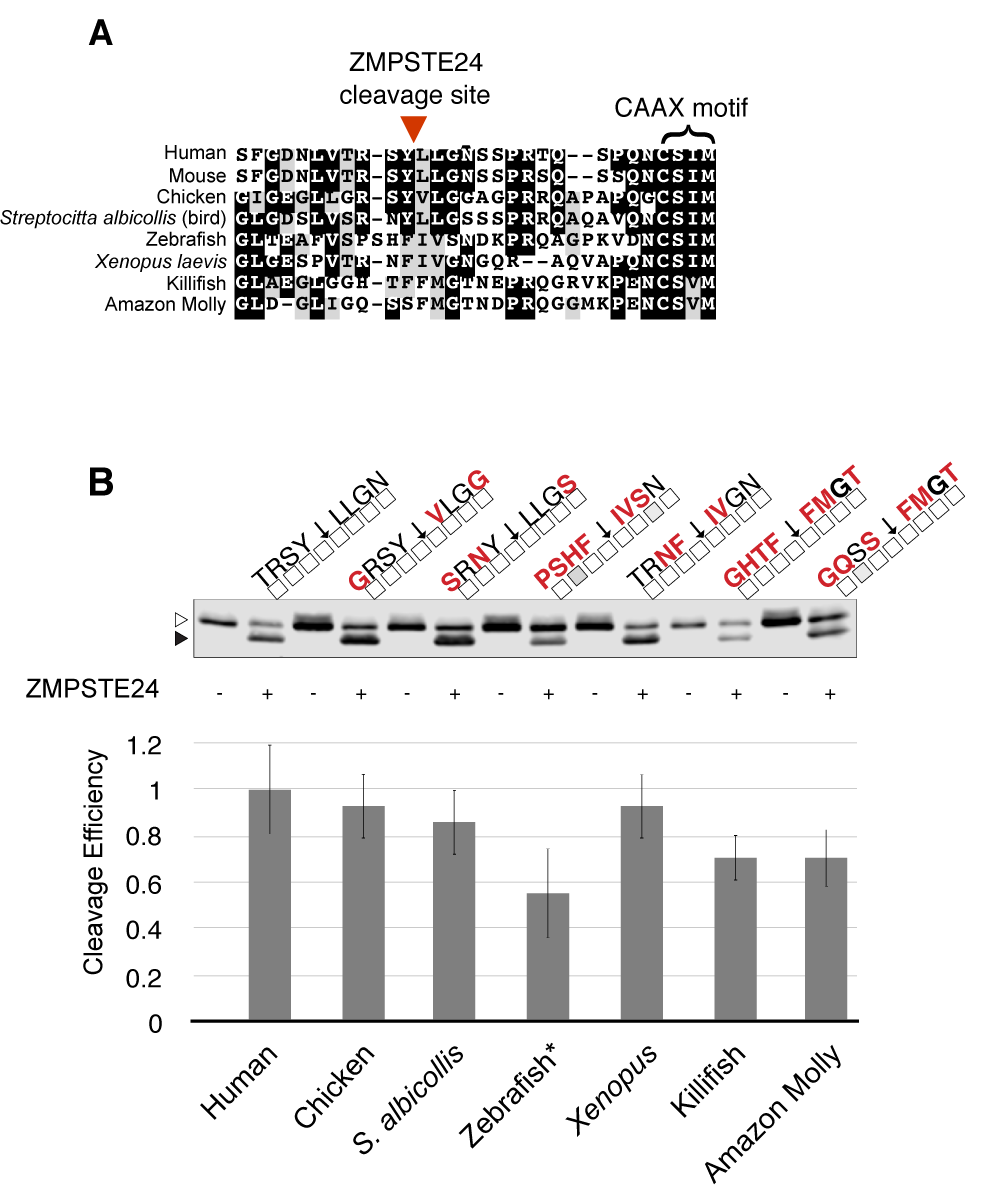
Multi-species analysis of ZMPSTE24-dependent cleavage of prelamin A. (A) Multi-species alignment of the amino acid sequence of the C-terminus of prelamin A. The inferred prelamin A cleavage site and CAAX motif are indicated. Black shading indicates identity and gray shading indicates similarity. (B) Western blots for inferred multi-species variants of the prelamin A cleavage site. Amino acid substitutions representing each species for P4-P4’ were made in plasmid pSM3393, as indicated above the western blot with differences from human prelamin A (left) highlighted in red. The white and gray boxes under the residues are in accordance with the figure 4 heatmap. Uncleaved prelamin A (empty arrowhead) and mature lamin A (filled arrowhead) are indicated. Each construct was assayed with (+) and without (-) ZMPSTE24 co-expression. Quantification was performed as in previous figures. White and gray boxes under residues are in accordance with heat map from Figure 4. The zebrafish sequence was the only divergent cleavage site with a statistically significant cleavage defect (p<0.05, Student’s T-test, two-tailed, unpaired, N=3), and is marked with an asterisk.

To ascertain the extent to which human ZMPSTE24 can cleave these divergent putative cleavage sites, we introduced amino acid substitutions in positions P4 through P4’ into our prelamin A expression construct to recapitulate the putative cleavage sites observed in the multi-species alignment. In the cleavage assay, all of the putative prelamin A cleavage sites were recognized and processed by ZMPSTE24, most with only modest differences relative to the human cleavage site TRSY↓LLGN (70-90%) (Figure 5B). The zebrafish sequence, with seven of the eight cleavage site residues substituted, was the only one with a statistically significant defect, although even in this case a substantial amount of cleavage (>50%) did occur. Thus, it appears that ZMPSTE24 can tolerate multiple substitutions at the cleavage site, so long as the changes conform to the rules defined by the heatmap data of Figure 4.

## Discussion

Here we have used a humanized yeast assay to determine the impact of amino acid substitutions in residues P4-P4’ surrounding the scissile bond of prelamin A for cleavage by ZMPSTE24. ZMPSTE24-dependent prelamin A processing *in vivo* is completely dependent on farnesylation (14,46-49). Farnesyl may be needed to deliver the substrate to membrane-embedded ZMPSTE24 and/or to engage a potential exosite inside or outside of the enzyme chamber. In the current work, the native yeast farnesyltransferase, (the Ram1/Ram2 heterodimer), along with the yeast CAAX processing enzymes Rce1 and Ste14 (11) generate a suitable farnesylated substrate for ZMPSTE24. This system has allowed us to interrogate a comprehensive library of farnesylated substrates for ZMPSTE24 cleavage.

Screening our collection of 152 cleavage site mutants in this *in vivo* assay yielded a heatmap of cleavage efficiency (Figure 4). In general, P1’ (Leu647) and P2’ (Leu648) were the most sensitive residues to perturbations, specifically alterations to non-hydrophobic mutations (Figures 3A and B). This is similar to, but distinct from, the related protease thermolysin which requires aliphatic or aromatic residues at P1’ but does not have a known preference at position P2’ (58). While substitutions at a few other positions of prelamin A (see below) diminish processing, almost none have as strong an effect as those at P1’ or P2’. At the outset of this study we knew that the P1’ position was critical for prelamin A cleavage, since the L647R mutation severely reduced prelamin A processing in cell-based experiments and a recently reported MAD-B patient (12,28,48,56,63). Here we found that no charged residues are allowable at P1’ and that hydrophobic residues are strongly preferred, except for the large aromatic residues tryptophan and tyrosine. Notably P2’ had similar requirements to P1’ although the negative effect of many substitutions at this position were generally diminished as compared to the same substitution at P1’.

To understand the potential impact of specific prelamin A mutations, we modeled the prelamin A tetrapeptide SY↓LL which contains the scissile bond and is comprised of residues P2-P2’, into the ZMPSTE24 cleavage site, based on our previously published structure of ZMPSTE24 and the CSIM tetrapeptide (PDB:2YPT) (31) (Figure 6A and B). The S1’ hydrophobic binding pocket is composed of Val332 and Leu283, and would not be expected to permit charged residues at P1’, just as we observed. The pocket is large enough to accommodate a large hydrophobic residue like phenylalanine, but not tryptophan. However, unlike L647F, L647Y could not be cleaved by ZMPSTE24. The reason probably lies in the difference between the abilities of tyrosine and phenylalanine to form hydrogen bonds. Hydrogen bonding by tyrosine with a nearby polar residue of ZMPSTE24 at the catalytic site (*e.g*., Glu415 or Arg465) could interfere with the catalytic reaction, which involves the attack of a water molecule on the carbonyl carbon of the scissile peptide bond. Indeed, substitution of Leu647 to any amino acid which can form a hydrogen bond resulted in poor cleavage efficiency.

**Figure 6:**
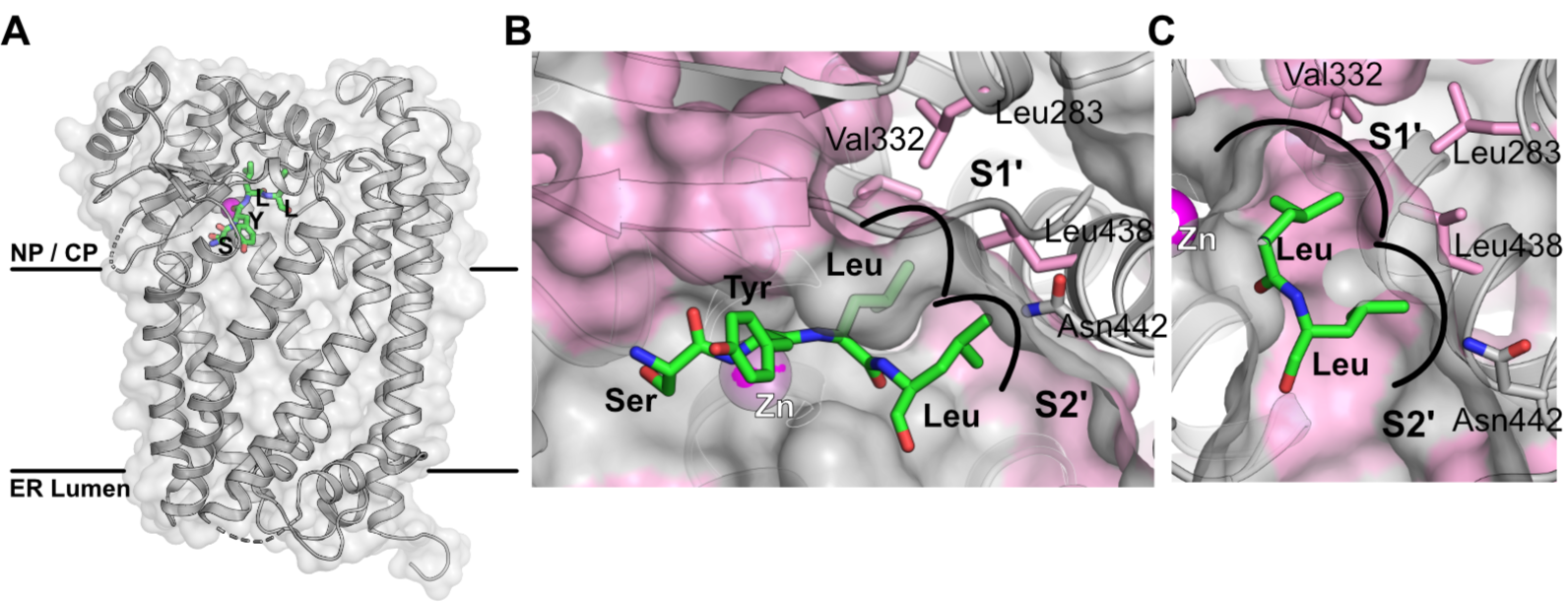
Model of the SYLL (P2-P2’ peptide) in the active site of ZMPSTE24. (A) The overall structure of ZMPSTE24, determined by X-ray crystallography, showing position of the active site Zn^2+^ (magenta) and with the prelamin A –SYLL substrate positioned in the proposed catalytic site, based on the co-crystal structure of ZMPSTE24 and the CSIM tetrapeptide (31). Substrate peptide backbone carbons are labeled green, the nucleoplasmic-cytoplasmic (labeled NP/CP) and ER lumen sides of the membrane in which ZMPSTE24 sits are indicated. (B) Zoom-in view of the active site, potential S1’ and S2’ binding pockets in the protease for P1’ and P2’ cleavage site residues of the substrate are annotated. (C) View of the S1’ and S2’ binding sites showing the hydrophobic residues forming the prelamin A peptide binding sites. Hydrophobic residues are in pink and labeled with three-letter amino acid residue code.

The predicted S2’ binding pocket has a hydrophobic surface formed by the sidechain of Leu438 (Figure 6B), consistent with our findings that at P2’ hydrophobic residues are also preferred. Poor cleavage of polar and charged residues in the P2’ position might be due to their hydrogen bonding with the nearby Asn442, resulting in misalignment of the substrate. In addition to the hydrophobic nature of the S1’ and S2’ binding pockets of ZMPSTE24, Leu647 and Leu648 on the prelamin A peptide contribute to the hydrophobic environment in this region. Consequently, a change of either residue to a more polar or charged sidechain is predicted to reduce the strength of the interaction of the adjacent sidechain. Similarly, a proline residue in this region would not only disrupt the interactions of that side chain, it would also misalign the sidechain of the adjacent residue, thus explaining the failure of cleavage with proline at either P1’ and P2’ sites.

The data presented here align with the few other studies performed to date on substrate specificity of ZMPSTE24/Ste24. Though not comprehensive, in a random mutagenesis screen for mutations that decreased yeast **a**-factor cleavage by Ste24 between residues T7 and A8 (at TAT↓ AAP), substitutions to the non-hydrophobic residues glycine or threonine at P1’ and proline at P2’ all blocked **a**-factor upstream processing (42). Purified Ste24 has been shown to correctly perform -AAX removal of a synthetic farnesylated **a**-factor 15-mer substrate using the native farnesylated **a**-factor CAAX motif f-CVIA (44), which conforms to the rule of hydrophobic residues at P1’ and P2’ (valine and isoleucine, respectively). Surprisingly, ZMPSTE24-dependent cleavage of the natural prelamin A CAAX sequence, f-CSIM, in the context of a farnesylated 29mer prelamin A peptide produced mostly a dipeptide, –IM, instead of the –SIM tripeptide expected for authentic –AAX cleavage (31,45). However, when the CAAX motif was changed to the Ras-derived sequence CAAX motif, farnesylated-CVLS, ZMPSTE24 produced a correctly –AAXed tripeptide, –VLS, highlighting the preference of the protease for hydrophobic residues (valine and leucine in this case) at the P1’ and P2’ positions (45).

ZMPSTE24 is able to recognize and cleave evolutionarily divergent P4-P4’ cleavage sites in prelamin A sequences obtained from several species (Figure 5). Although prelamin A processing has only been directly demonstrated for human, mouse, and chicken homologs, the strong similarity observed in alignments of the C-terminal portion of prelamin A for numerous vertebrate sequences suggests ZMPSTE24 processing is likely to be a shared feature among all species. None of the residues flanking these putative cleavage sites (Figure 5) violate the rules defined by our heatmap. All have “allowable” hydrophobic residues at the inferred P1’ and P2’ sites, and indeed all were cleaved when placed into the context of prelamin A at a level 50-100% that of WT. While the rules established in this study are unlikely to be specific enough to be predictive of possible substrates (i.e. to propose a strict consensus motif), it is notable that human ZMPSTE24 could recognize and cleave several divergent substrate sequences that conform to the rules indicated by the heatmap.

Although substrate farnesylation increases both efficiency and fidelity of cleavage, it is not an absolute requirement for ZMPSTE24/Ste24-dependent processing *in vitro* (31,45). Along these lines, yeast Ste24 has been shown to cleave non-farnesylated substrates unrelated in sequence to prelamin A or **a**-factor, including an M16A protease reporter and an insulin B-chain. Mass spectrometry analysis of cleavage products revealed that the P1’ and P2’ sites chosen within these substrates for cleavage by Ste24 *in vitro* were hydrophobic in nature (50). While translocon clogger substrates have not yet been evaluated in terms of their ZMPSTE24-dependent cleavage products, it would be of interest to analyze these products by mass spectrometry.

We also observed in this study that the arginine at position P3 is sensitive to substitution. Negatively charged residues, hydrophobic residues, and particularly tryptophan, all diminished cleavage. This residue is of particular interest because of the association of variant R644C with diverse human disease phenotypes including cardiovascular disease, muscular dystrophy, and skeletal abnormalities. This variant shows incomplete penetrance in families (61). It has also been detected as a particularly common variant in the ExAC database (64). Neither we nor others have observed a prelamin A cleavage defect in patient cells bearing this mutation (63,65), although we did observe a partial cleavage defect for R644C in a mammalian expression system where the amount of available substrate is difficult to control (63). In the present study, R644C shows approximately 80% cleavage efficiency compared to WT (Figure 2B; Figure 4). The impact of R644C on prelamin A processing, and whether the associated disease phenotypes are due to an accumulation of prelamin A in cells of patients harboring this mutation, remains unclear. R644H has also been reported to be associated with disease, but disease causality has not been proven (66). In our study this R644H amino acid change did not affect processing. Overall, the findings for changes in residue R644 (position P3) are ambiguous in contrast to the L647R (position P1’) mutations, where we see a severe processing defect and cells from a MAD-B R644C patient have been shown to exhibit a strong accumulation of prelamin A (62).

There are limitations in this study. The assay we used here reports steady-state cleavage efficiency in whole-cell lysates and does not measure the rates of processing of prelamin A variants by ZMPSTE24. Also, this assay does not ascertain the exact cleavage site, although gel migration of the cleaved product is consistent with cleavage occurring at or near the predicted cleavage site (SY↓LL). Mutation of a protease cleavage recognition sequences can result in shifting the cleavage site, and we cannot exclude this possibility for our mutants that show successful cleavage. In any case, because of the importance of ZMPSTE24 cleavage in human health and disease, gaining a better understanding of how this enzyme selects its substrates can contribute to our understanding both of progeroid diseases and also the normal physiological process of aging.

## Experimental Procedures

### Yeast Strains Used in this Study

The yeast strain SM4826 (*ste24Δ::kanMX his3 leu2 met15 ura3*) contains a deletion of the *STE24* gene and is from the BY4741 deletion collection (67). Strain SM6303 (*ste24::KanMX TRP1::P_PGK1_-His_10_-HA_3_-ZMPSTE24*) was used as the recipient strain for plasmids containing mutant forms prelamin A and analyzing their cleavage efficiency. It is derived from strain SM4826 and contains two copies of human *ZMPSTE24* (codon optimized for yeast (68)) integrated into the genome at the *TRP1* locus (59).

### *LMNA* Plasmids and Mutagenesis

All *LMNA* mutants in this study are variants of plasmid pSM3393, a centromeric *HIS3* plasmid derived from pRS313 (69). In pSM3393 (shown in Fig. 1), the human prelamin A C-terminal tail (LMNA_CT_; corresponding to amino acids 431-664 of prelamin A) is N-terminally tagged with *His*_*10*_-*myc*_*3*_ and is expressed from the *PRC1* promoter (59). The collection of 152 *LMNA* substitution mutants and 6 multispecies variants assayed in this study were generated by QuikChange™ mutagenesis (Stratagene, San Diego, CA) with degenerate oligonucleotides or, when necessary, by individual mutagenic PCR reactions combined with NEBuilder^®^ HiFi Assembly (New England Biolabs, Ipswich, MA). Plasmid pSM3393 was the template for all reactions. For QuikChange™ mutagenesis, antiparallel oligonucleotides were designed to contain a degenerate codon, NNK (N represents G, T, A, or C and K represents G or T), at the target codon. Mutagenic QuikChange™ reactions were performed on template plasmids that had been deleted for the target codon, to reduce the incidence of wild-type amplification products. QuikChange™ reactions were *Dpn*I-treated, transformed into NEB^®^ 5-alpha competent *E. coli* cells, and selected on carbenicillin-containing LB media. Plasmid DNA was prepared from individual colonies and individual mutations were identified and confirmed by Sanger sequencing.

For each position (P4-P4’; prelamin A residues 643-650), nearly all amino acid substitutions were generated by the degenerate oligonucleotide approach. However, when a particular mutation was missing after degenerate oligonucleotide mutagenesis, antiparallel QuikChange™ primers were individually designed and used with outside primers to generate mutation-containing PCR products for *in vitro* assembly with the pSM3393 backbone. The combination of degenerate oligonucleotide mutagenesis and individual PCR-based mutagenesis yielded the complete collection of 152 lamin A cleavage site mutants. The multispecies variants analyzed in Fig. 6 were made by QuikChange mutagenesis, using antiparallel oligonucleotides designed to contain the multiple substitutions of each species’ putative lamin A cleavage site.

The WT *LMNA*_*CT*_ and 152 *LMNA*_*CT*_ variants on pSM3393 were individually transformed into the yeast strain SM6303, (or in some cases SM4826), by standard lithium acetate protocols, selected on minimal SC-His plates, and transformants were re-streaked to isolate single colonies.

### Prelamin A Cleavage Assay

Overnight cultures were grown at 30°C to saturation in minimal medium (0.67% yeast nitrogen base, 0.5% ammonium sulfate, 2% glucose, supplemented with amino acids), back diluted 1:1000, and cells were grown for 14-16 hours to an OD_600_ of approximately 1.0. Cells (1.0 OD_600_ cell equivalents) were pelleted, washed with water, treated with 0.1 N NaOH, and lysed in SDS-PAGE sample buffer at 65° C for 10 minutes (70). Lysates were vortexed, pelleted, and supernatants (0.24 OD_600_ cell equivalents per lane) were resolved on 10% SDS polyacrylamide gels for 90 minutes at 150V. Proteins were transferred to nitrocellulose membranes using the Trans-Blot^®^ Turbo™ system (Bio-Rad Laboratories, Hercules, CA) and blocked with Western Blocking Reagent (Roche, Indianapolis, IN).

Lamin A proteins were detected by probing with mouse anti-Myc antibodies (clone 4A6, Millipore cat #05-724; 1:10,000 dilution) decorated with goat anti-mouse secondary IRDye 680RD antibodies (LI-COR, Lincoln, NE). The loading control hexokinase was detected using and rabbit anti-hexokinase (1:200,000 dilution) antibodies decorated with goat anti-rabbit secondary IRDye 800CW (LI-COR) antibodies. Western blots were visualized using the Odyssey^®^ imaging platform (LI-COR) and bands were quantified using Image Studio™ software (LI-COR). Cleavage efficiency was calculated by dividing mature lamin A signal by total lamin A signal (processed + unprocessed) and normalized to WT processing efficiency, which was set to 1.0. Cleavage assays were performed as 3 independent biological replicates. Blots were re-probed using rat anti-HA (clone 3F10, Roche cat #11867423001; 1:10,000 dilution) to detect ZMPSTE24 and visualized using goat anti-rat IRDye 680RD.

### Multispecies Alignment

Protein sequences for the species studied were acquired from UniProt (UniProt Consortium, 2017) and aligned using ClustalX (71). Putative cleavage sites for divergent species were inferred from the alignment and introduced into pSM3393 via mutagenic PCR and assembled as described above. Resulting constructs were transformed into yeast strains SM6303 and SM4826 and assayed for prelamin A cleavage as described above.

## Acknowledgments

We thank Mark Dumont (University of Rochester) for providing a yeast codon-optimized version of human *ZMPSTE24*. We thank Kaitlin Wood and Khurts Shilagardi, members of the Michaelis lab for comments on the manuscript and Ashley Pike in the Carpenter lab for help with Figure 6.

## Funding and additional information

This work was funded by a National Institute of Health (NIH) grant to SM (5R35GM127073) and a Medical Research Council (MRC) grant (MR/L017458/1) to EPC. LN and EPC are funded by the Structural Genomics Consortium (SGC). The SGC is a registered charity (number 1097737) that receives funds from AbbVie, Bayer Pharma AG, Boehringer Ingelheim, the Canada Foundation for Innovation, Genome Canada, Janssen, Lilly Canada, Merck KGaA, Merck & Co., Novartis, the Ontario Ministry of Economic Development and Innovation, Pfizer, São Paulo Research Foundation-FAPESP and Takeda, as well as the Innovative Medicines Initiative Joint Undertaking ULTRA-DD grant 115766 and the Wellcome Trust (106169/Z/14/Z).

